# A panel of phenotypically and genotypically diverse bioluminescent:fluorescent *Trypanosoma cruzi* strains as a resource for Chagas disease research

**DOI:** 10.1101/2024.03.27.586912

**Authors:** Francisco Olmo, Shiromani Jayawardhana, Archie A. Khan, Harry C. Langston, Amanda Fortes Francisco, Richard L. Atherton, Alex I. Ward, Martin C. Taylor, John M. Kelly, Michael D. Lewis

## Abstract

Chagas disease is caused by *Trypanosoma cruzi*, a protozoan parasite that displays considerable genetic diversity. Infections result in a range of pathological outcomes, and different strains can exhibit a wide spectrum of anti-parasitic drug tolerance. The genetic determinants of infectivity, virulence and therapeutic susceptibility remain largely unknown. As experimental tools to address these issues, we have generated a panel of bioluminescent:fluorescent parasite strains that cover the diversity of the *T. cruzi* species. These reporters allow spatio-temporal infection dynamics in murine models to be monitored in a non-invasive manner by *in vivo* imaging, provide a capability to detect rare infection foci at single-cell resolution, and represent a valuable resource for investigating virulence and host:parasite interactions at a mechanistic level. Importantly, these parasite reporter strains can also contribute to the Chagas disease drug screening cascade by ensuring that candidate compounds have pan-species *in vivo* activity prior to being advanced into clinical testing. The parasite strains described in this paper are available on request.

**AUTHOR SUMMARY:** Chagas disease results from infection with the protozoan parasite *Trypanosoma cruzi* and is a major public health problem throughout Latin America. *T. cruzi* is a genetically diverse species and infection can result in a wide range of pathological outcomes, mainly associated with the heart and/or digestive tract. Research on Chagas disease, ranging from fundamental biology to drug development, has been greatly aided by the availability of genetically modified parasite reporter strains that express bioluminescent:fluorescent fusion proteins. In combination with mouse models and imaging technology, these strains allow infections to be monitored in real-time, with high sensitivity, and infection foci to be visualised at single-cell resolution. Here, we describe an extensive panel of bioluminescent and fluorescent strains that cover the diversity of the *T. cruzi* species. These reporter strains, that are available on request, should have wide utility in many areas of Chagas disease research. In particular, as part of the drug development screening programme, they can be used to ensure that candidate compounds have *in vivo* activity across the species prior to being advanced into clinical testing.

## INTRODUCTION

Chagas disease is the most serious parasitic infection in the Americas, with 6-7 million people infected with causative agent, the insect-transmitted protozoan *Trypanosoma cruzi* [1]. Infections with this obligate intracellular parasite are typically life-long, and in 30-40% of cases, result in chronic cardiac and/or digestive tract pathology, although this can take decades to become symptomatic [2, 3]. Globally, Chagas disease is the leading cause of infectious cardiomyopathy. The front-line drugs used to treat infections are the nitroheterocyclic agents benznidazole and nifurtimox [4, 5]. However, therapeutic failure is common, with toxicity and long administration periods (>60 days) often leading to treatment cessation [6, 7]. In addition, the wide diversity of the *T. cruzi* species is associated with considerable variations in the level of therapeutic susceptibility, even amongst strains that are closely related at an evolutionary level [8].

Anti-Chagasic drug research is challenging. Progress has been restricted by several factors, including the long time periods over which symptoms develop, the wide spectrum of disease pathology, and the limited accuracy of the diagnostic tools used to determine curative outcomes. For example, during the chronic stage of disease, highly sensitive PCR-based methodologies can be confounded by the low and fluctuating parasite load in blood, and the highly focal nature of infections within deep tissue sites [9]. In addition, robust biomarkers for cure are lacking. As a result, predictive animal models have assumed an important role in the drug development pipeline, and have been central to research on disease pathogenesis and immune mechanisms. *In vivo* bioluminescence imaging of murine infections has been established as an experimental tool applicable to Chagas disease research [10–12]. However, the sensitivity required to monitor infection dynamics throughout the course of the disease was not achieved until parasites were engineered to express a codon-optimised, red-shifted luciferase [13–16]. The enhanced qualities of this reporter stem from the improved tissue penetration of longer wavelength light towards the red-end of the spectrum. This is due to reduced light scattering and decreased absorbance (haemoglobin is the major chromophore in tissue). Imaging applications have been further extended by linking a fluorescent protein encoding sequence (mNeonGreen) in-frame with the luciferase gene, to generate parasites that express a bioluminescent:fluorescent fusion protein [17]. This has enabled *in vivo* foci of *T. cruzi* infections to be visualised at single cell resolution, even during the chronic stage of the disease when the parasite burden is extremely low [18–21].

The genetically diverse *T. cruzi* species has been sub-divided into six major lineages (Discrete Typing Units – DTUs; TcI – TcVI) [22]. However, factors such as virulence, tissue tropism, and drug-sensitivity display both inter- and intra-lineage variation [8, 14, 23, 24], and the genetic basis of these phenotypic traits is poorly understood. Recently, progress in Chagas disease drug development has been accelerated by the formation of multidisciplinary research consortia that have combined expertise from the academic, not-for-profit and commercial sectors [25–29]. These projects have highlighted a need to better capture the extent of parasite diversity within the screening cascade. Here, we describe a wide range of parasite reporter strains that cover all six DTUs, and which display a spectrum of infection profiles. The availability of these strains will provide a means of ensuring that candidate compounds have cross-species *in vivo* efficacy before advancement into clinical trials. In addition, they represent a valuable research resource for identifying the genetic determinants of infectivity and virulence.

## Results and Discussion

The bioluminescent *T. cruzi* CL Brener (TcVI) strain has been widely used for *in vivo* imaging in murine models of Chagas disease, particularly in the context of drug testing [eg. 9, 16, 25, 27 - 29]. This parasite line (*T. cruzi* CLBR-Luc) was generated by transfection with vector pTRIX2-RE9h (S1A Fig) [13, 30], which targeted the *PpyRE9h* red-shifted luciferase gene [31] to a high expressing ribosomal DNA (rDNA) locus. To generate reporter strains that are both bioluminescent and fluorescent, the red-shifted luciferase gene can be replaced with a *PpyRe9h-mNeonGreen* fusion sequence (S1B Fig) [17]. These strains can be further modified by replacement of the *mNeonGreen* sequence with *mScarlet* [17] (S2 Fig; Materials and Methods), to generate red fluorescent variants. Parasite lines that express dual reporters have infection profiles indistinguishable from the parental bioluminescent clones (S3 Fig), and a limit of detection of <20 parasites when infected mice are assessed by *ex vivo* imaging [19].

To produce a panel of reporter strains that encompasses the broad diversity of the *T. cruzi* species, we selected a range of isolates covering each of the major human infectious lineages (DTU I - VI) (Table 1). These 19 strains are from wide geographical origins within South America, and are derived from both insect and mammalian hosts (predominantly human). They encompass an 8-fold variation in benznidazole sensitivity (4 – 32 μM), and a ∼3-fold variation in doubling time when cultured as epimastigotes (18 – 48 hours). There is no correlation between replication rate and benznidazole sensitivity (Table 1). Each of these strains was transfected with the construct pTRIX2-RE9h (S1A Fig), selected with G418 and cloned transformants then assessed to confirm bioluminescence (Materials and Methods). Many of the parasite lines were also modified further to express bioluminescent:fluorescent fusion proteins following integration of genes encoding mNeonGreen and/or mScarlet (Table 2; Materials and Methods). Selected parasite clones were then used to generate infectious *in vitro* trypomastigotes and to infect CB17 SCID mice, an immunodeficient line that lacks functional lymphocytes [32]. The majority of these reporter parasite strains underwent exponential growth in SCIDs, such that humane endpoints were reached within 28 days (Fig 1, S4 Fig, Table 2). However, some parasite strains displayed a less virulent phenotype in these immunodeficient mice, particularly Pot7a (DTU II), Rita (DTU II), X10610 (DTU IV) and BUG2148 (DTU V).

**Fig 1.**
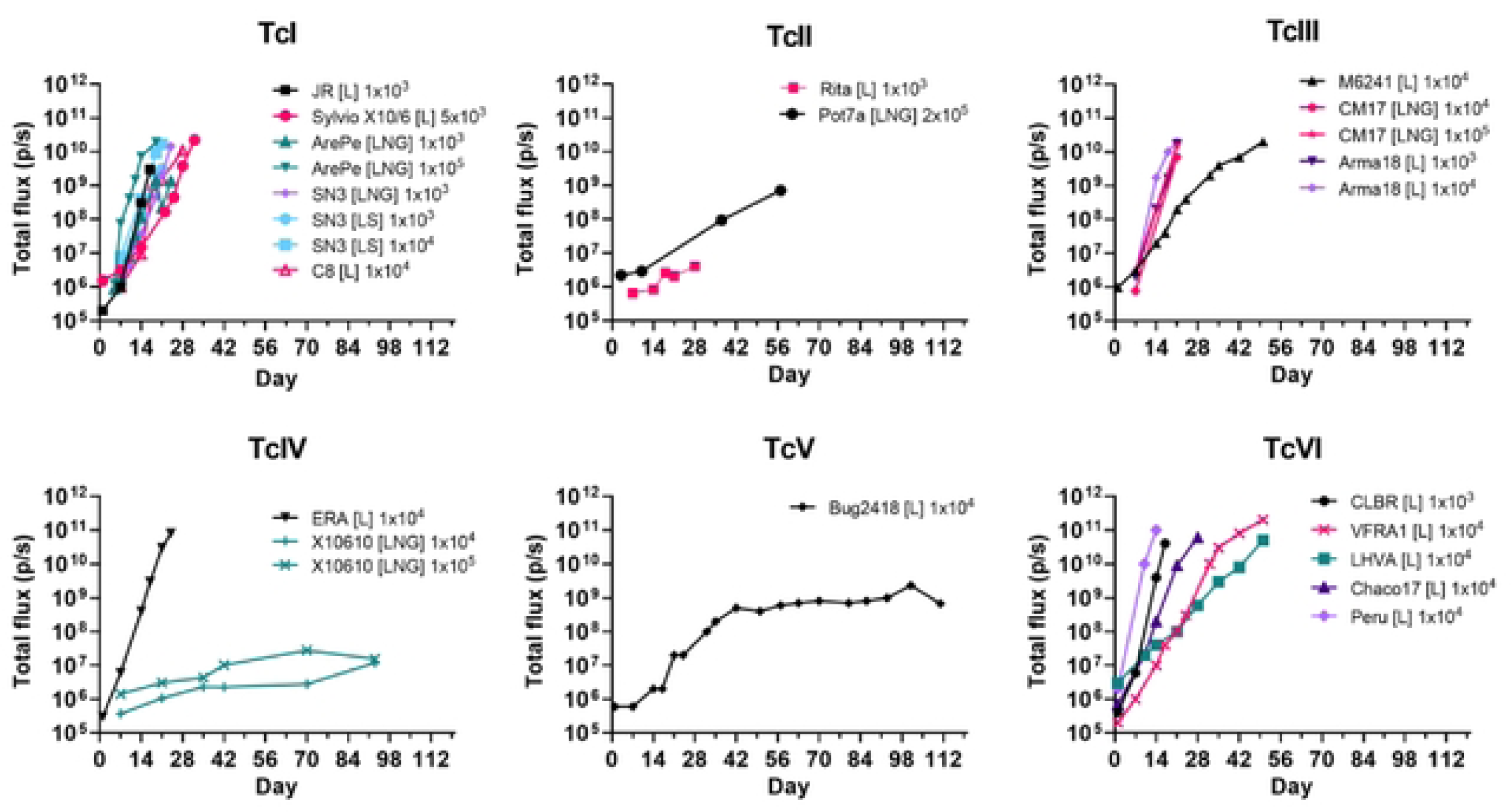
Growth profile of selected *T. cruzi* reporter strains in severe combined immunodeficient (SCID) mice. CB17 SCID mice were infected with a range of *T. cruzi* strains that expressed red-shifted luciferase (L), or as indicated, fusion proteins also containing mNeonGreen (LNG) or mScarlet (LS) fluorescent components (Table 2) (Materials and Methods). Inoculations were by the i.p. route, with the number of tissue culture trypomastigotes, as indicated. The mice were monitored regularly by bioluminescence imaging and the total flux (p/s) from ventral view images recorded (Materials and Methods). The number of days post-infection is indicated. Mice were euthanized at humane end-points by exsanguination under terminal anaesthesia.

We next investigated infection dynamics in BALB/c and C3H/HeN mice, immunocompetent strains that are widely used in *T. cruzi* research, including drug efficacy studies. In total, 23 mouse:parasite strain combinations were assessed. These encompassed each of the major *T. cruzi* lineages, and parasite strains expressing either the red-shifted luciferase reporter, or bioluminescent:fluorescent fusion proteins (mNeonGreen or mScarlet) (Table 2).

With BALB/c mice, the infection profiles of 8 *T. cruzi* strains covering 4 DTUs (I, III, V and VI) were monitored in detail. The *in vivo* bioluminescence of 6 of these strains followed a similar trend, with the parasite burden peaking 14 – 21 days post-infection, followed by a steady reduction as the infection transitioned from the acute to the chronic stage (Fig 2A). After day 50, in most cases, bioluminescence-inferred parasite burdens reached a dynamic steady-state characterised by transient infection foci which appeared and disappeared over time. This type of profile has been observed previously in BALB/c infections with the bioluminescent CLBR (DTU VI) and JR (DTU I) strains [13, 14]. The SN3 strain (DTU I) exhibited a lower acute stage peak than observed with other infections, but in the chronic stage, the bioluminescent profile was similar to infection with other strains, including the highly dynamic nature of infection foci. In contrast, infections with the CM17 strain (DTU III), which resulted in the highest parasite burden during the acute phase (Fig 2A), became barely detectable as the chronic stage proceeded (>100 days). In one case, with the BUG2148 strain, bioluminescence was not detected beyond day 4 post-infection.

**Fig 2.**
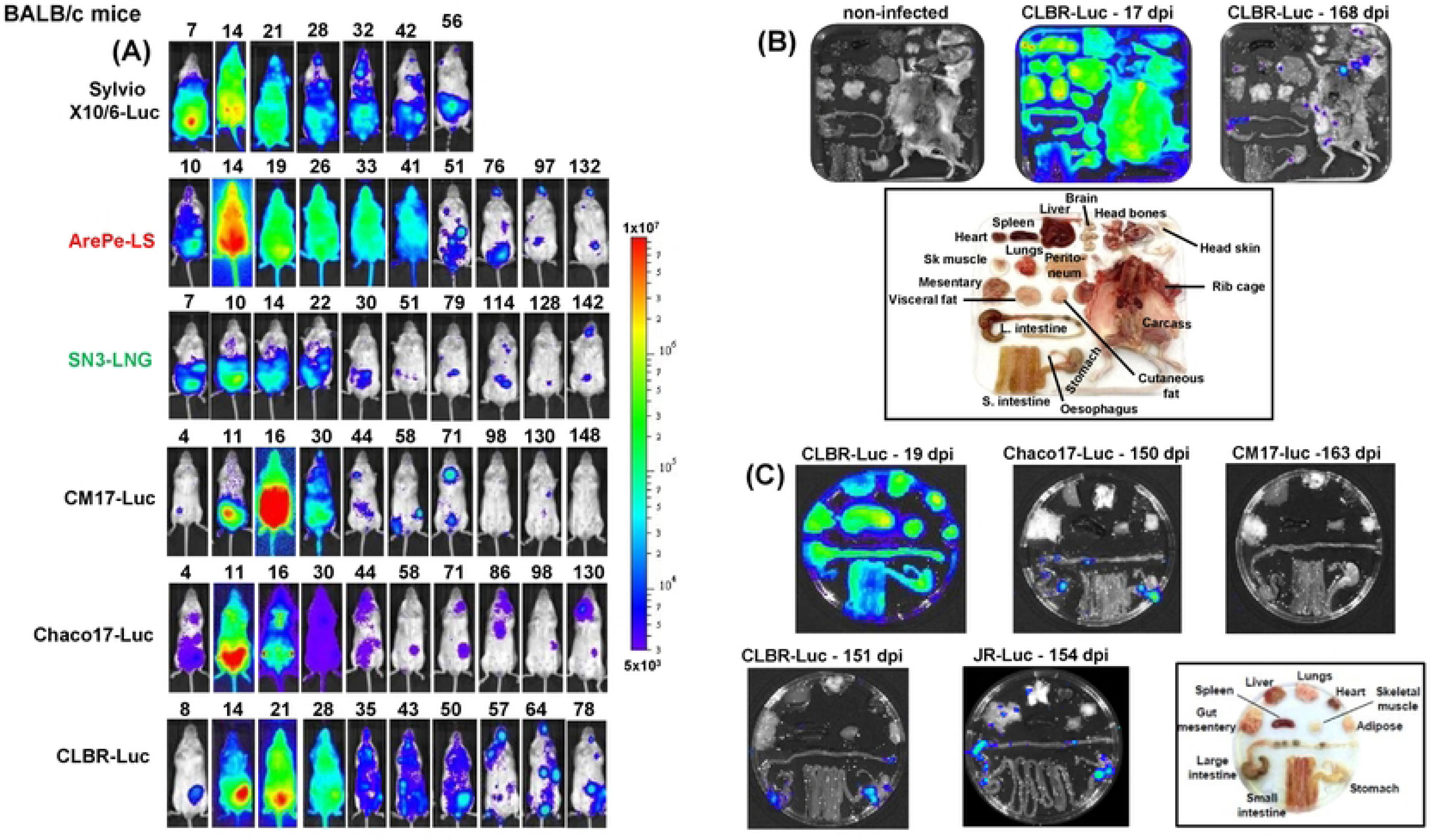
Bioluminescence imaging of BALB/c mice infected with a range of *T. cruzi* strains. (A) Dynamics of infection. BALB/c mice were infected with *T. cruzi* reporter strains by the i.p. route and monitored regularly by *in vivo* imaging until they had transitioned to the chronic stage (Materials and Methods). Ventral images are shown and the number of days post-infection indicated. -Luc; strains expressing the red-shifted luciferase PpyRE9h. -LS, -LNG; strains expressing luciferase:mScarlet and luciferase:mNeonGreen fusion proteins, respectively. Further details on parasite strains are provided in Table 1 and 2. (B) Assessment of the parasite burden in tissues and organs of *T. cruzi* CLBR-Luc infected mice during the acute (17 days post-infection; dpi) and chronic (168 dpi) stages. Mice were injected with D-luciferin prior to necropsy, during which the organs and tissues were harvested, arranged as shown (inset), and examined by *ex vivo* imaging (Materials and Methods). (C) Assessment of parasite burden in tissues and organs of BALB/c mice during chronic stage infections with a range of strains as indicated. Organs from a *T. cruzi* CLBR-Luc infected mouse during the acute stage (19 dpi) are shown for comparison. The heat-maps are on a log_10_ scale and indicate intensity of bioluminescence from low (blue) to high (red); the minimum and maximum radiances for the pseudocolour scale are indicated. This scale was used for both *in vivo* and *ex vivo* images.

In the acute stage, where examined, parasites were widely disseminated, with all organs and tissues highly infected (Fig 2B and 2C). When BALB/c infections progressed to the chronic stage, parasites became mainly restricted to distinct foci in the musculature of the GI tract (predominantly in the colon and/or stomach) and skin, with infections of other organs and tissues being more intermittent (Fig 2C). This type of infection profile has been reported previously with the CLBR and JR strains [13, 14, 19]. On the basis of the chronic tissue-specific infection profiles for the strains analysed, it appears that the GI tract serves as an immunologically permissive reservoir for *T. cruzi* in most BALB/c mouse infections.

Nine *T. cruzi* strains across five DTUs (I, II, III, IV, VI) have been tested in C3H/HeN mice. The extent of infection tended to be more variable (Fig 3A). The highest parasite burdens resulted from infections with the Peru (DTU VI), CLBR and JR strains, with the peak of the acute stage occurring later than in BALB/c mice, and stretching over a longer period (days 14 - 35 post-infection). Other strains were less infectious, particularly Pot7a, where bioluminescence fell below the threshold of detection from day 35 onwards. It should be noted that the light brown fur of C3H/HeN mice reduces the sensitivity of *in vivo* imaging compared to the white-furred BALB/c mice, due to differential absorbance of the bioluminescence signal. When assessed by *ex vivo* imaging, in the chronic stage, parasite strains were generally more widely disseminated in tissues and organs than observed with BALB/c mice, and it is less clear whether there are major differences between sites in their permissiveness to long-term infection (Fig 3B and 3C) [14, 20].

**Fig 3.**
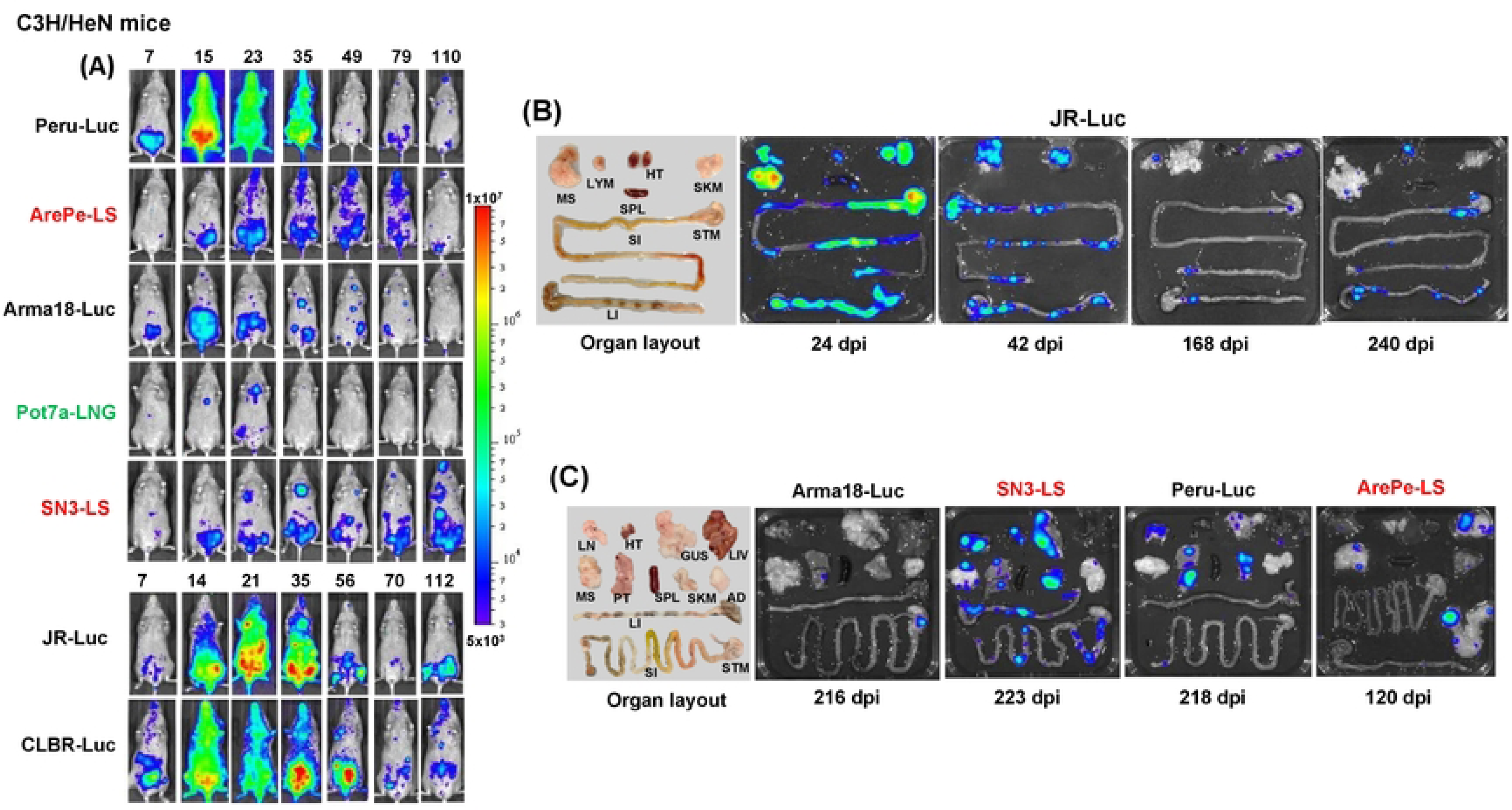
Bioluminescence imaging of C3H/HeN mice infected with a variety of *T. cruzi* reporter strains. (A) Dynamics of infection. C3H/HeN mice were infected as described (Materials and Methods), and monitored by *in vivo* imaging (ventral view shown) at the days post-infection, as indicated. The bioluminescence scale and abbreviated strain names are as in Fig 1 (see Table 2 for further details). (B) *Ex vivo* imaging of organs and tissues harvested from *T. cruzi* JR-Luc infected mice over the course of infection (Materials and Methods). (C) *Ex vivo* images of organs and tissues from mice chronically infected with a range of reporter strains. Days post-infection (dpi) are indicated. The same heat-map scale indicating intensity of bioluminescence was used for both *in vivo* and *ex vivo* images. Organ layout is shown (left) using the following abbreviations: lymph nodes – LYM, Lungs - LN, gut mesenteric tissue - MS, heart - HT, spleen - SPL, skeletal muscle - SKM, stomach - STM, small intestine - SI, large intestine – LI, genito-urinary system – GUS, liver – LIV, peritoneum – PT, and adipose tissue – AD.

Parasites expressing bioluminescent:fluorescent proteins (Table 2) also provide an opportunity to examine chronic infections at single-cell resolution and to assess their immunological and tissue microenvironments (Materials and Methods) [19, 33]. Previously, this revealed that some infected host cells avoid detection by the immune system, whereas others induce extensive immune cell infiltration into the locality [21]. Using *T. cruzi* CLBR PpyRE9h:mNeonGreen strain infections of C3H/HeN mice (Table 2), we can show that the tissue-level microenvironment is not a determinant of this differential response. In colonic smooth muscle, infection foci targeted by T cell infiltration can co-exist with other closely localised (100 - 200 μM) parasite “nests”, that have equal or greater infection loads, but do not trigger a simultaneous immune response (Fig 4A). Bioluminescence-guided targeting followed by fluorescence microscopy of individual infected cells can also been used to visualise skin localised parasites in the dermal layer [19] and to study enteric nervous system pathology in the context of local *T. cruzi* infection foci [20] (Fig 4B, C, D). These types of study, together with the expanded panel of transgenic parasites therefore provide a platform to study immune evasion and tissue pathogenesis in the context of parasite genetic diversity. This should contribute to a better understanding of why *T. cruzi* infections are associated with such highly variable clinical outcomes.

**Fig 4.**
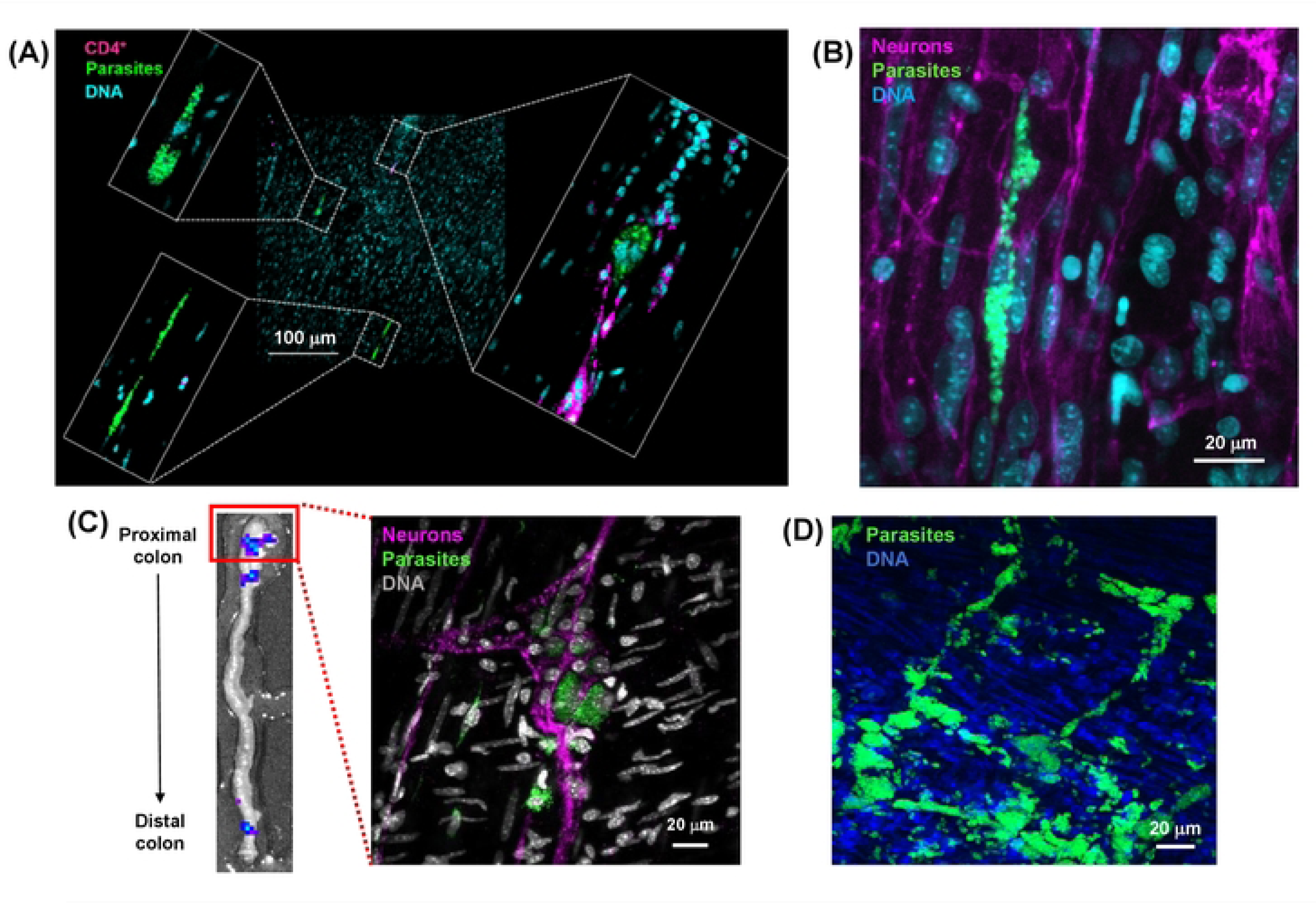
The use of bioluminescent:fluorescent *T. cruzi* reporter strains to monitor tissue microenvironment in the colon during infection. (A) Differential recruitment of T cells to closely localised *T*. *cruzi* CLBR-Luc:NeonGreen chronic stage (121 dpi) infection foci in a C3H/HeN mouse colon. Smooth muscle wall sections were prepared from colonic whole mounts, following bioluminescence-guided excision, and screened using a Zeiss LSM880 confocal microscope to localise fluorescent infection foci [18–20] (Materials and Methods). The image expansions show three closely localised infection foci, only one of which has triggered recruitment of T cells. Parasites, green (fluorescence); DNA, cyan (Hoechst 33342); CD4^+^ T cells, magenta. (B) Chronic stage (230 dpi) *T. cruzi* CLBR-Luc:NeonGreen infections of colonic smooth muscle in the context of local enteric neurons. C3H/HeN tissue sections were prepared as above. Neurons, magenta (TuJ1+). (C) A whole mount colon tissue section from a C3H/HeN mouse infected with *T. cruzi* JR-Luc:NeonGreen imaged by fluorescence microscopy, showing parasites (green) in the myenteric nerve plexus (magenta) (139 dpi). The inset (left) shows the proximal colon bioluminescence image that was used to guide tissue excision and fluorescent imaging of infection foci at single cell resolution (Materials and Methods). (D) Whole mount colonic tissue obtained from an immunocompromised SCID mouse during a fulminant infection with *T. cruzi* CLBR-Luc:NeonGreen (21 dpi) at the level of the myenteric nerve plexus (unlabelled).

### Concluding Remarks

The extent of *T. cruzi* genetic diversity has been a major barrier to progress in Chagas disease research, and the impact of parasite genetics on infection dynamics, pathogenesis and drug susceptibility remain unresolved questions. There are advantages to focusing and co-ordinating research around one or a few favoured laboratory strains, but this must be balanced against the need to generate data that can usefully reflect the true diversity of *T. cruzi* infections. The panel of reporter strains described here therefore represents a valuable resource that will be widely applicable and should improve the translational value of data generated using *in vitro* and *in vivo* experimental infection models. For example, it will allow mixed infections, which are common in humans, to be studied in detail, both *in vitro* (S5 Fig) and *in vivo*, and provide a framework within the Chagas disease drug development pipeline for testing the pan-species *in vivo* efficacy of candidate compounds. In combination with other advances, such as the generation of mice humanised for drug-metabolising enzymes expressed by the cytochrome P450 superfamily [34], this should increase the predictive power of *in vivo* pre-clinical testing.

## Materials and Methods

### Parasite culturing and generation of genetically engineered strains

All *T. cruzi* strains used in this study are described in Table 1. They were genotyped to the DTU level using a previously published triple locus AFLP/RFLP assay [35]. Epimastigotes were cultured at 28°C in supplemented RPMI-1640 medium as described previously [14]. Metacyclic trypomastigotes were generated following transfer of epimastigotes to Graces transformation medium, in some cases supplemented with filter-sterilised triatomine intestinal homogenate [14, 36]. Tissue culture trypomastigotes were obtained from infected MA104 cells grown at 37°C using minimal Eagle medium supplemented with 5% heat-inactivated foetal bovine serum (FBS), or from infected L6 rat skeletal myoblasts grown in RPMI-1640 medium, supplemented with 10% heat-inactivated FBS [14].

The generation of bioluminescent *T. cruzi* CL Brener parasites expressing the red-shifted luciferase gene *PpyRE9h* [31] has been described previously [13]. Bioluminescence was conferred on other strains by integrating *PpyRE9h* into the rDNA locus following transfection with a 6.0 kb DNA fragment derived by Aat II/Asc I digestion of vector pTRIX2-RE9h (GenBank accession #PP332292) (S1A Fig), and selection of transformants with G418 [13, 30]. To generate CL Brener parasites expressing bioluminescent:fluorescent fusion proteins, the red-shifted luciferase gene was replaced with a *LucRe9h-mNeonGreen* fusion sequence to produce the parasite line *T*. *cruzi* CLBR-Luc::NeonGreen [17]. Transfection with a 7.0 kb Sac I/AscI fragment derived from construct pTRIX-LucRe9h-mNeonGreen (GenBank accession #OR258373) (S1B Fig), and selection with hygromycin allowed the generation of green fluorescent variants from across the DTU spectrum (Table 2). For parasite lines already containing a red-shifted luciferase gene in a high expressing rDNA locus, a 5.3 kb SbfI/AscI fragment can be used to target the fluorescent component into this transcriptionally active site. Red fluorescent variants of the fusion protein were first generated in the CL Brener strain by replacement of the *LucRE9h* gene with the dual reporter *LucRE9h-mScarlet* sequence using the T7 RNA polymerase/cas9 system, and selection with blasticidin [17]. Transfection of other strains with a 5.35 kb PCR product derived from this cell line (GenBank accession #PP332293) (S2A Fig) allowed the generation of a series of red fluorescent variants (Table 2). Subsequently, we have produced construct pTREX2-LucRe9h-mScarlet (GenBank accession #PP333636) (S2B Fig), which contains a 7.0 kb SacI/AscI fragment that can be used to generate red fluorescent variants following transfection and selection with hygromycin.

### Drug sensitivity assays

For activity assays, logarithmically growing *T. cruzi* epimastigotes were sub-cultured at 2.5 x 10^5^/ml into 96-well plates at a range of benznidazole concentrations, and incubated for 4 days. Resazurin was added and the plates incubated for a further 2 days, then read using a BMG FLUOstar Omega plate reader. These times were modified, as required, to account for slow growing strains. Results were analyzed using GraphPad Prism to determine the drug concentrations that inhibited growth by 50% (EC_50_). Experiments were performed in triplicate.

### Ethics statement

Animal studies were performed under UK Home Office project licences (PPL 70/8207 and PPL P9AEE04E4), and ratified by the LSHTM Animal Welfare and Ethical Review Board (AWERB). Procedures were carried out in accordance with the UK Animals (Scientific Procedures) Act 1986.

### Mice and infections

BALB/c, C3H/HeN and CB17 SCID mice were purchased from Charles River (UK). They were housed under specific pathogen-free conditions in individually ventilated cages. They experienced a 12-hour light/dark cycle and had access to food and water *ad libitum*. Female mice, ranging in age from 6–12 weeks were used for infections. SCID mice were typically infected with 1 × 10^4^ *in vitro* derived tissue culture trypomastigotes in 0.2 ml PBS via i.p. injection. BALB/c and C3H/HeN mice were also infected by i.p injection, in most cases with 10^3^ or 10^4^ bloodstream form trypomastigotes derived from parasitaemic SCID mouse blood. Most SCID mice developed fulminant infections (Fig 1, S4 Fig) and were euthanized at, or before, humane end-points by ex-sanguination under terminal anaesthesia (euthatal or dolethal; 15 μl/g body weight) [37].

### Bioluminescence imaging

For *in vivo* imaging, mice were injected i.p with 150 mg/kg d-luciferin in DPBS (Dulbecco’s Phosphate Buffered Saline), and anaesthetized by 2.5% (v/v) gaseous isoflurane in oxygen [13, 37]. To assess bioluminescence, they were placed in an IVIS Lumina II or Spectrum system (Revvity, Hopkinton, MA, USA) and images acquired using Living Image 4.7.2 software. Exposure times varied from 30 seconds to 5 minutes, depending on signal intensity. Mice were then revived and returned to their cages. For *ex vivo* imaging, mice were injected with 150 mg/kg d-luciferin i.p. as above, and then sacrificed by lethal i.p. injection with dolethal 5 minutes later. They were perfused with 10 ml of 0.3 mg/ml luciferin in DPBS via the heart, the organs and tissues were removed, transferred to a round or square Petri dish in a standardized arrangement, soaked in 0.3 mg/ml luciferin, and imaged using the IVIS [14].

### Fluorescence imaging

Histological sections were generated after bioluminescence-guided excision of infection foci from murine tissues [17, 37]. Biopsy specimens were incubated in 95% EtOH at 4°C overnight and then washed in 100% EtOH (4 x 10 minutes), followed by xylene (2 x 10 minutes). Samples were embedded by placing in melted paraffin wax, which was allowed to set. The embedded pieces were cut into histological sections (5 – 20 μm) using a microtome, the sections were then processed as previously described [37], mounted in Vectashield, and imaged using a Zeiss LSM880 confocal microscope.

For whole mounts, colonic muscularis samples were prepared as previously described [19–21]. Briefly, colon pieces were opened, stretched and pinned on Sylgard 184 plates, then fixed in 4% paraformaldehyde, after which the mucosal layer was removed using fine forceps. Tissues were washed with PBS, permeabilised (1% Triton X-100 in PBS), and blocked (10% sheep serum, 1% Triton X-100 in PBS). Anti-CD4 rat antibodies (Abcam) at 1:500 were used to assess C4^+^ T cell recruitment, and neurons were labelled with rabbit anti-tubulin beta 3 (TuJ-1) IgG at 1:500 (Biolegend), in PBS, with 0.5% Triton X-100 for 48 hours at 4°C. Tissues were washed, incubated with secondary goat anti-rat IgG (Invitrogen) or goat anti-rabbit-AF633 IgG (ThermoFisher), respectively, in PBS containing 0.5% Triton X-100 for 2 hours and counterstained with Hoechst 33342 (1 µg/ml) at room temperature. Whole mounts were examined and imaged by confocal microscopy as above.

---

**S1 Fig.**
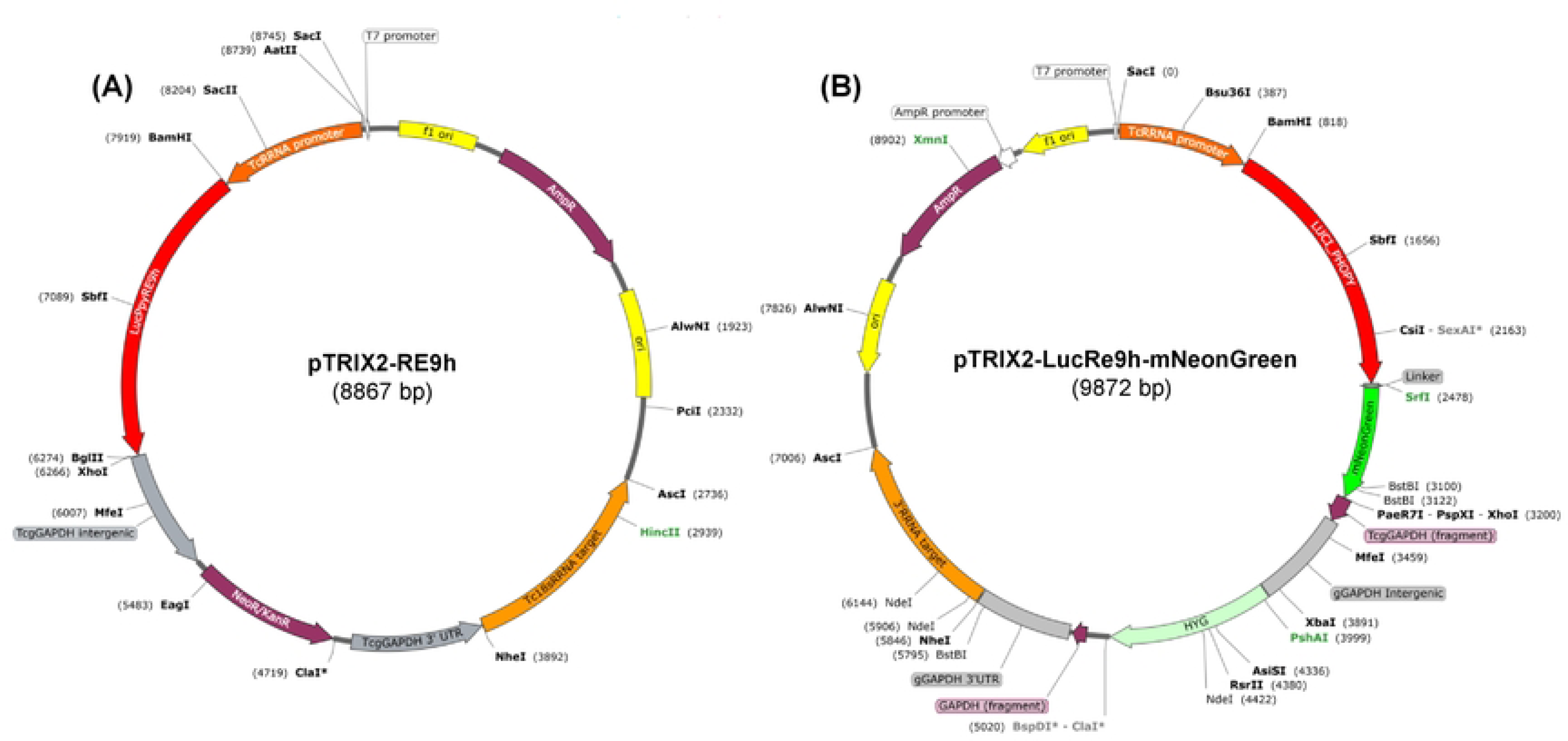
Constructs used to generate *T. cruzi* bioluminescent:green fluorescent reporter strains. (A) The red-shifted luciferase gene *PpyRE9h* [31] can be targeted to *T. cruzi* rDNA loci after transfection with a 6 kb fragment produced by AatII/AscI digestion of construct pTRIX2-RE9h and selection with G418 [13, 30]. (B) The PpyRE9h:mNeonGreen fusion sequence [17] can be similarly targeted following transfection with a 7.0 kb SacI/AscI fragment fragment from construct pTRIX2-LucRe9h-mNeonGreen, and selection with hygromycin. Construct sequences are available on GenBank (Materials and Methods).

**S2 Fig.**
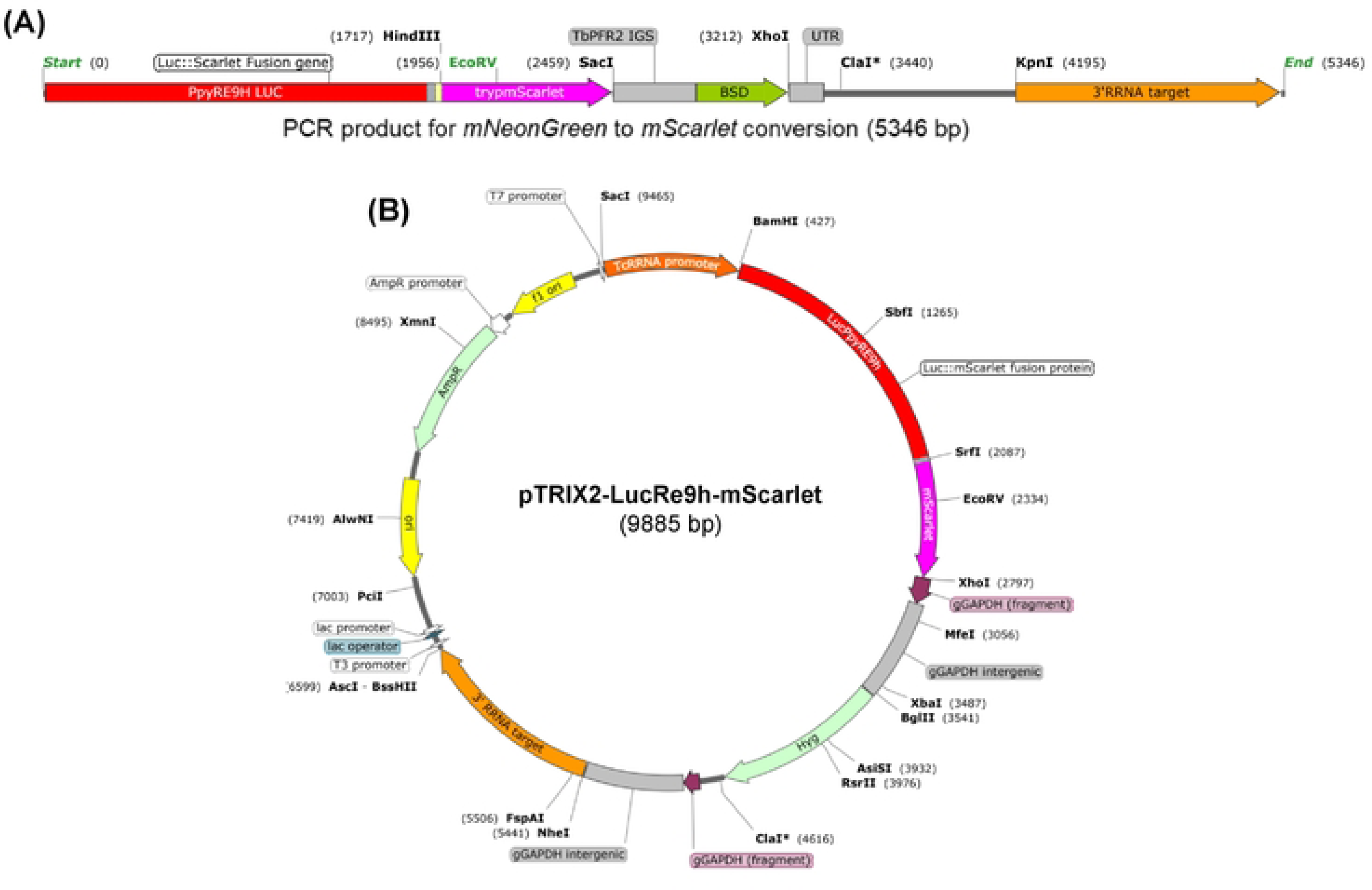
Constructs used to generate *T. cruzi* bioluminescent:red fluorescent reporter strains. (A) Map of the 5.35 kb PCR fragment used to convert the fluorescent component of the dual reporter protein from mNeonGreen to mScarlet with the blasticidin S deaminase gene (BSD) as the selectable marker [17]. (B) Construct pTRIX2-LucRe9h-mScarlet which contains a 7.0 kb SacI/AscI fragment that can be used to transfect *T. cruzi* and generate red fluorescent variants following selection with hygromycin. Sequences of fragment and construct are available on GenBank (Materials and Methods).

**S3 Fig.**
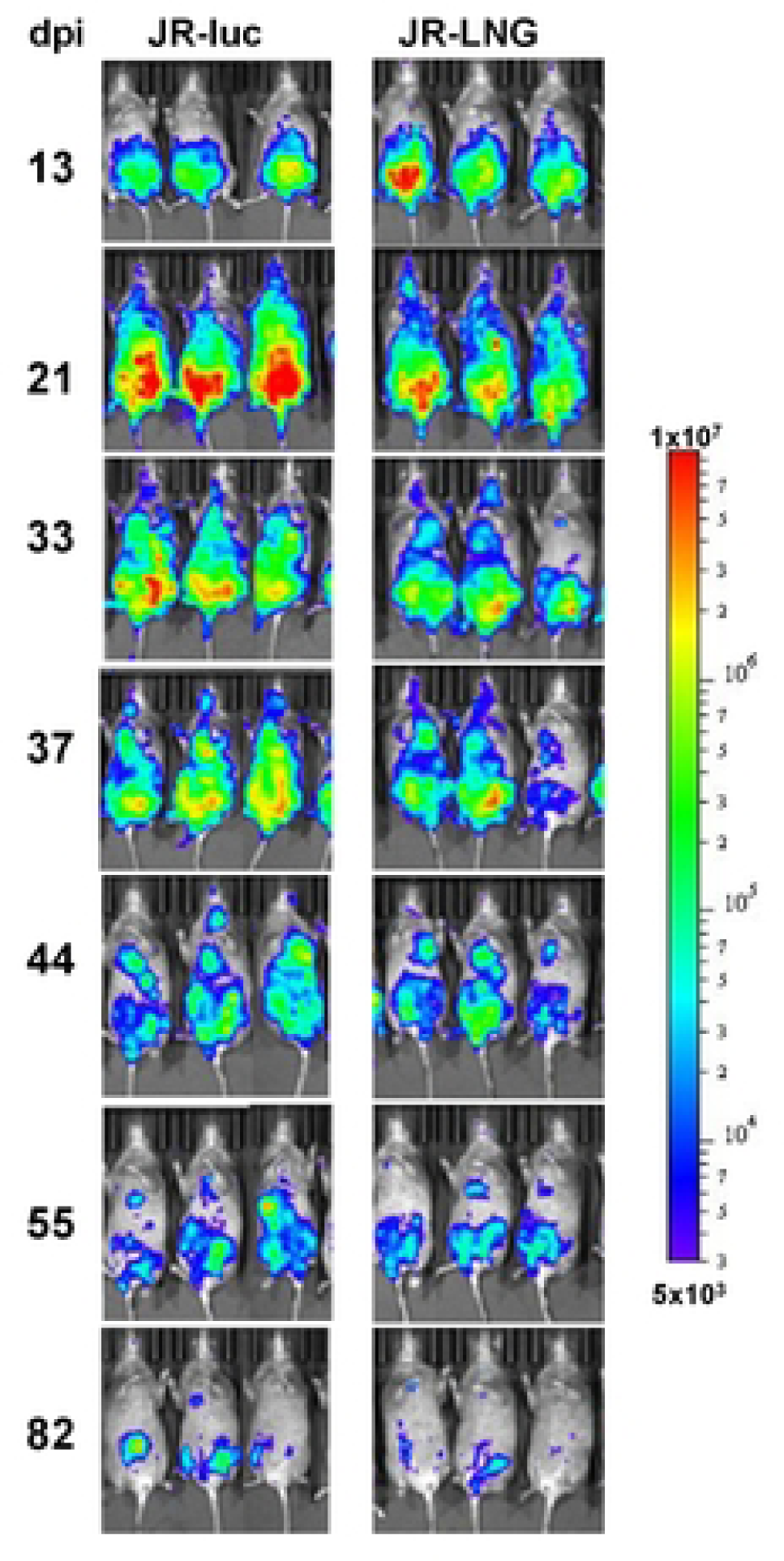
Monitoring infection dynamics of *T. cruzi* strain JR expressing different reporter proteins. C3H/HeN mice were injected i.p. with 5 x 10^4^ trypomastigotes expressing either **the** red-shifted luciferase (JR-Luc) or the luciferase:mNeonGreen fusion protein (JR-LNG) (Materials and Methods). They were monitored by *in vivo* imaging, at the days indicated post-infection (dpi). The heat-map indicates the intensity of bioluminescence.

**S4 Fig.**
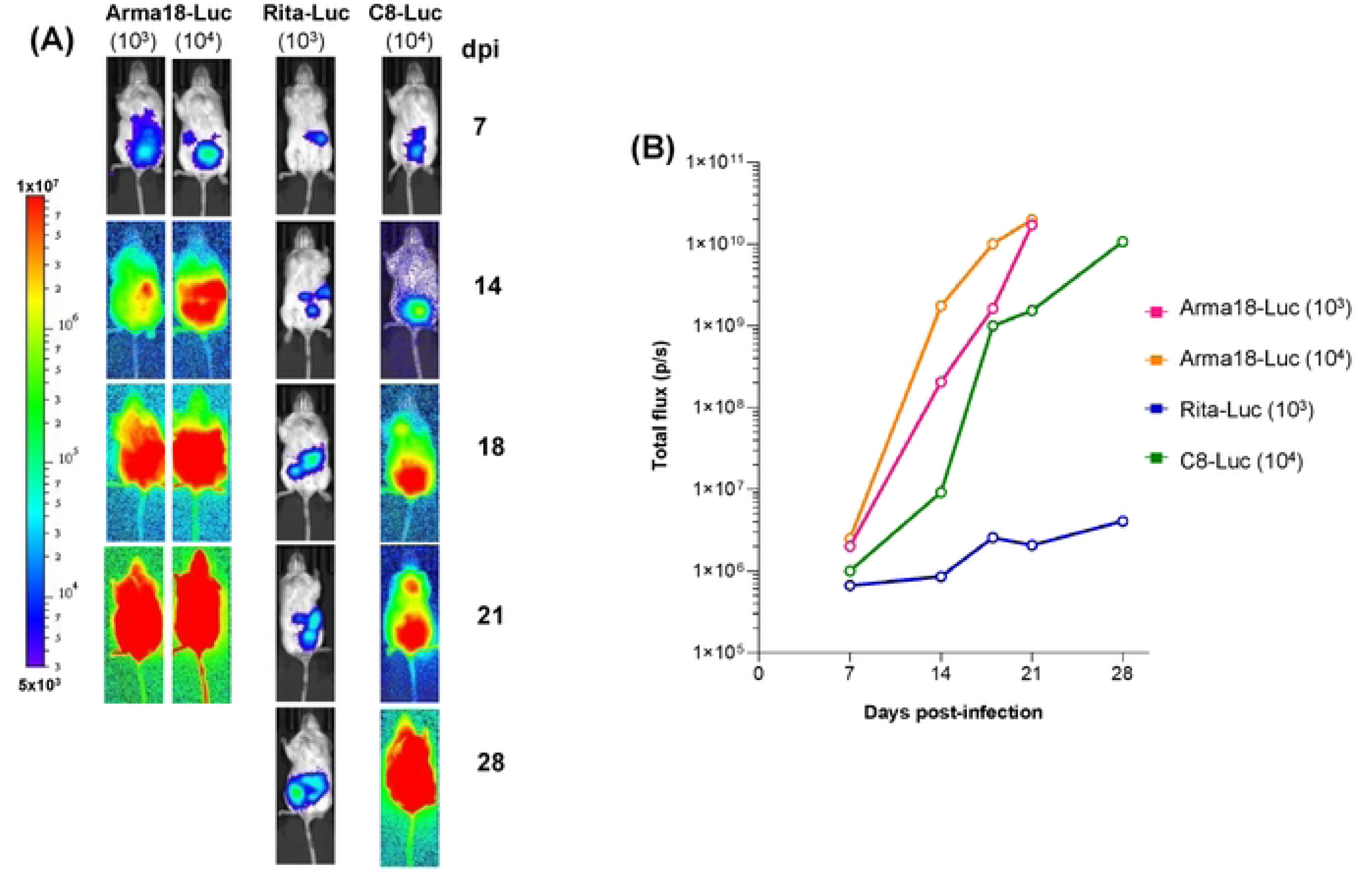
Differential infection dynamics of *T. cruzi* strains in immunodeficient SCID mice. Mice were infected i.p. with bioluminescent tissue culture trypomastigotes (numbers shown in brackets) and monitored by *in vivo* imaging (Materials and Methods) at the days indicated post-infection (dpi). The heat-map indicates the intensity of bioluminescence. (B) Total bioluminescence flux derived by ventral imaging at the days indicated.

**S5 Fig.**
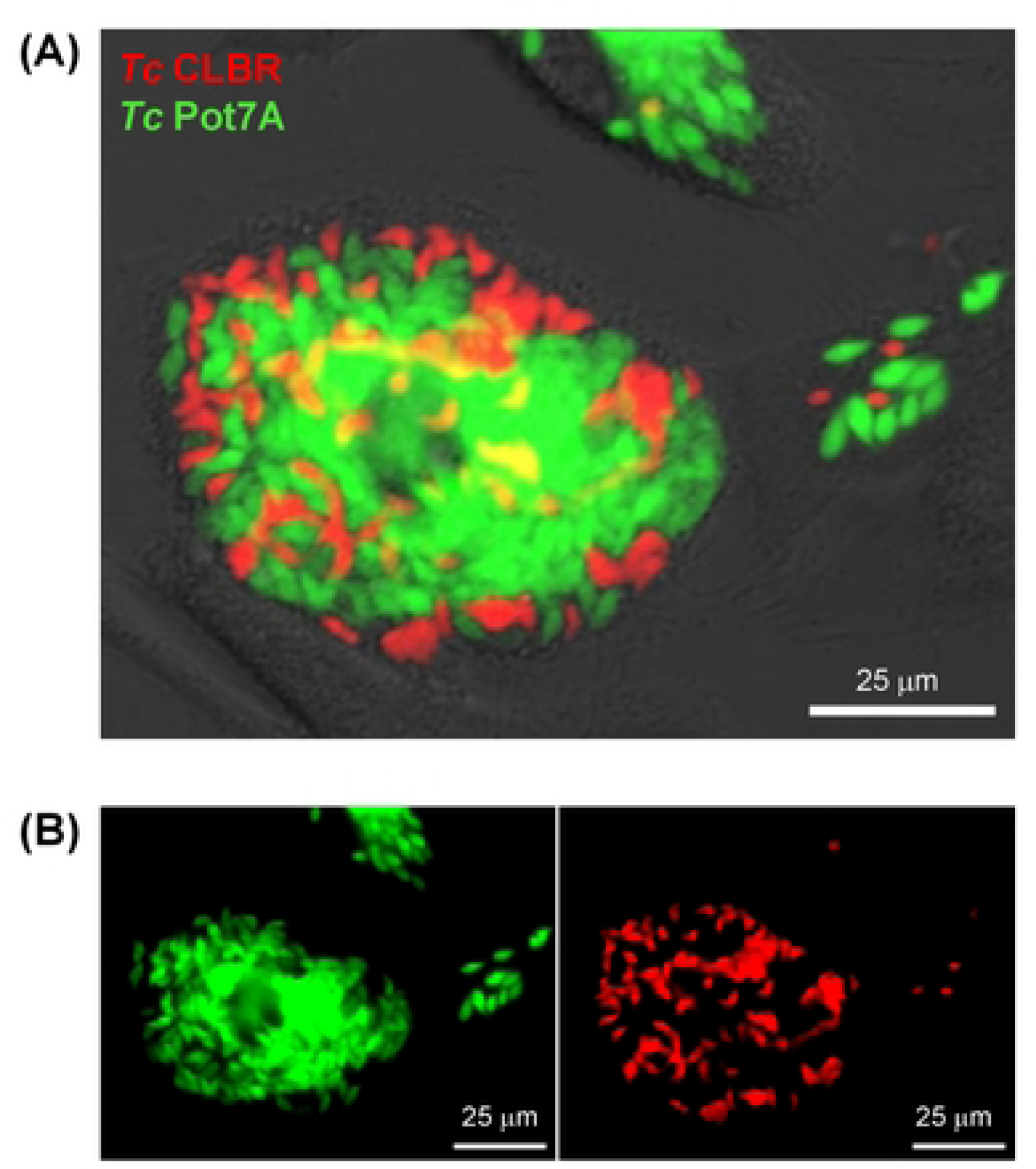
The use of distinct dual reporter strains to assess the impact of co-infection. Human foreskin fibroblast (HFF) cells were infected with *T. cruzi* Pot7a-Luc:NeonGreen (*Tc* Pot7a – green) and 5 days later with *T*. *cruzi* CLBR-Luc:mScarlet (*Tc* CLBR – red). (A) Image taken with a Nikon Ti-2 E inverted microscope using red and green filters, 5 days after the second infection. (B) Same image captured with red or green filters. Epifluorescence real-time microscopy can also be used to track co-infections.

## Acknowledgements

This work was supported by UK Medical Research Council (MRC) grants MR/T015969/1 to J.M.K. and MR/R021430/1 to M.D.L., and funding from the Drugs for Neglected Diseases initiative (DNDi). DNDi received financial support from: Department for International Development (DFID), UK; Federal Ministry of Education and Research (BMBF) through KfW, Germany; and Médecins sans Frontières (MSF) International. AIW was in receipt of an MRC LID (DTP) Studentship (MR/N013638/1).

The funders had no role in study design, data collection and analysis, decision to publish, or preparation of the manuscript.

